# Reinforcement of speciation by behavioural and reproductive barriers in the *Drosophila bipectinata* species complex

**DOI:** 10.1101/2025.10.31.684612

**Authors:** M Manjunath, C S Damini, V Shakunthala

## Abstract

Speciation mainly depends on different mechanisms involved in reproductive isolation; Understanding such barriers that consistently develop to avoid the fitness cost through reinforcement until the speciation continuum completes helps clarify the role of pre-mating and post-mating isolating mechanisms. The *Drosophila* bipectinata species complex, comprising four closely related taxa, provides a natural system for studying the interplay of behavioural, sensory, and genetic mechanisms underlying reproductive isolation. In this study, a comprehensive characterization was undertaken on twelve different types of sterile F1 hybrid males among the crosses between four species of *Drosophila* bipectinata species complex which are experiencing reproductive isolation. Comparing the mating latency and copulation duration among the pure species and hybrids, showed a significant increase in the mating latency of hybrids but the duration of copulation among certain hybrids of *Drosophila* bipectinata, *Drosophila* parabipectinata and *Drosophila* malerkotliana showed no significant reduction in the copulation time and emphasized the strong influence of X chromosome inheritance of hybrid genome. Experimental ablation of antennae demonstrated that male olfactory input is indispensable for heterospecific mating success, consistent with the role of pheromone detection in conspecific recognition, while tarsi ablation revealed striking sex-specific asymmetry: male tarsi removal increased heterospecific success, whereas female tarsi removal abolished mating entirely, highlighting the role of gustatory receptors in female mate choice. Testis dissections further confirmed that post-meiotic defects in hybrids, including failure of spermatid elongation and individualization, these findings demonstrates that reproductive isolation in the *D. bipectinata* species complex arises from the collective action of multiple barriers, prezygotic isolation mediated by pheromonal and gustatory divergence, and postzygotic sterility driven by spermatogenic failure. Our study underscores how behavioural and physiological divergence interact to reinforce reproductive barriers, providing a comprehensive framework for understanding speciation in *Drosophila*.

## INTRODCUTION

Speciation is a continuous process through the gradual accumulation of behavioural, developmental, and ecological incompatibilities in a diverging population (Mayr 1963, Coyne and Orr 2004). Speciation usually occurs as a result of physical or biological barriers that develops among the incipient species, which ultimately prevents interbreeding(Mayr 1942). Biological barriers among recently diverged taxa can happen due to many different incompatibilities where they have developed genetic differences between males and females before or after the initiation of copulation(Svensson et al., 2007). Physically incapable to copulate or behavioural rejections prior to copulation, which might happen due to differences in male mating signals or pheromonal indifferences among the heterospecific females results in pre-copulatory reproductive isolation(Kozak et al. 2009). Achieving complete reproductive isolation (RI) and inhibiting the gene flow requires the accumulation of multiple barriers to evolve step by step gradually which will take over thousands to millions of generations. Considering pre-copulatory isolation as one of the major reproductive barriers, the duration of copulation indirectly specifies the probabilities of successful mating or transfer of sperm among the given conspecific or heterospecific pairs(Sagga et al. 2011, Manier et al. 2013). In one of the studies of the *Drosophila simulans* clade, heterospecific matings between *Drosophila simulans* females and *Drosophila sechelia* males the copulation duration is considerably shorter than conspecific matings and further examination revealed that very few sperm has transferred(Price et al. 2001). Even if the copulation duration is long enough to complete the sperm transfer when the reproductive isolation is still in its continuum and has not yet achieved complete speciation, post mating pre zygotic isolating mechanisms undertake the spotlight and complete the isolation, in such a condition when females have multiple matings with different males successful fertilization depends on the post-copulatory mechanisms such as cryptic female choice (Eberhard 1996, Firman et al. 2017) and sperm competition (Parker 1970). There are examples of sperm dumping’s by females and poor sperm storage when an female is mated to heterospecific males which negatively affects the heterospecific sperm (Matute and Coyne 2010). In *Drosophila*, behavioural barriers such as divergence in cuticular hydrocarbons (CHCs), pheromone profiles, and species-specific courtship songs act as major prezygotic filters (Coyne et al., 1994; Higgie & Blows, 2007; Yew & Chung, 2017). These sensory signals are processed through olfactory and gustatory receptor pathways, which mediate male recognition and female acceptance during mating. Even minor changes in pheromone composition or receptor sensitivity can reduce heterospecific mating, reinforcing RI in sympatric populations (Smadja & Butlin, 2009; Billeter et al., 2012).

In natural hybrid zones, post-copulatory mechanisms interact in such a manner that competition arises between heterospecific and conspecific sperms, in which heterospecific ejaculation could be less suited to the female ethology to fertilize and to lay eggs or to produce a healthy progeny, ending up with a reduced fecundity in crosses with heterospecific males and permitting conspecific sperm to carry through the majority of fertilization (Howard 1999, Howard et al. 2008). However, in laboratory conditions, these type of reproductive barriers has been well studied (Peterson et al. 2011, Larson et al. 2012, Turissini et al. 2017, Poikela et al. 2019) but the relative frequencies between conspecific and heterospecific matings are unknown for majority of the incipient species and it is relatively unclear in characterizing the specific steps through the simultaneous development of pre-mating and Pre-Mating Post Zygotic(PMPZ) barriers. This interaction between reduced hybrid fitness and prezygotic mate discrimination is the basis of reinforcement, whereby natural selection favors stronger assortative mating in sympatric populations (Servedio & Noor, 2003; Matute, 2010). Evidence from multiple *Drosophila* clades demonstrates that reinforcement can rapidly strengthen premating barriers through divergence in pheromones, mating latency, and copulation duration (Noor, 1995; Yukilevich, 2012; Hopkins, 2013).

Apparently, the common outcome prior to complete reproductive isolation is hybrid male sterility. Most of the theoretical and empirical studies have shown that sterility in interspecies male hybrids is the result of tenuous disruption during sperm development (spermatogenesis), typically most of the investigations reveal post-meiotic problems such as asynchrony in the spermatid development and its failure to differentiate into mature sperm cells, sperm unindividualisation, aggregation of cellular debris in sperm bundles, and formation of undifferentiated interconnected spermatids resulting in immotile sperms (Haerty and Singh 2006, Gomes and Civetta 2014). Sometimes hybrid males having single motile sperm will be considered fertile, but the movement of sperm alone can’t ensure fertility because there are examples of hybrid male sterility having elusive physiological defects even though the sperms are motile enough but incapable to fertilize the eggs (Civetta and Gaudreau 2015).

Hybrid males of *bipectinata* species complex are completely sterile bidirectionally and the preliminary works on the morphology of testes and seminal vesicles (Singh and Mishra 2005) have shown that the absence of motile sperm as the cause of hybrid male sterility among this species complex, which shows that the perturbance of spermatogenesis in hybrid males is causing the sterility but the particular stages at which spermatogenesis is disturbed is pretty much unclear. The *bipectinata* complex consists of four species, *Drosophila bipectinata* (Duda 1923), *Drosophila malerkotliana* (Parshad and Paika 1964), *Drosophila parabipectinata* (Bock 1971) and *Drosophila pseudoananassae* (Bock 1971). Females of all the species are indistinguishable, males can be identified by abdominal tip pigmentation and sex combs. Despite extensive work on the genetic basis of hybrid male sterility, far fewer studies have simultaneously quantified both behavioral barriers and postmating defects in the same species complex. Understanding how mating latency, copulation duration, and spermatogenic abnormalities interact offers a more complete picture of how multilayered isolation evolves. The *D. bipectinata* complex, with its four closely related species and complete bidirectional hybrid male sterility, provides an ideal system to dissect the contributions of precopulatory and postcopulatory prezygotic mechanisms in parallel.

## MATERIALS AND METHODS

### *Drosophila* stocks and Hybridization regime

The *Drosophila* species used in this study were *Drosophila bipectinata* Pune, *Drosophila parabipectinata* Mysore, *Drosophila malerkotliana malerkotliana* Raichur (all from India) and *Drosophila pseudoananassae nigrens* Brunei Island, Brunei. All of the stocks were obtained from BHU, Varanasi, India. The stocks were reared in bottles containing yeast agar medium and maintained at 23^0^-24^0^ C in 12hrs:12hrs light-dark cycle with 70% humidity of laboratory conditions. Approximately 40 flies (males and females in equal number) were transferred to fresh culture bottles for every 10 days, also the progenies were transferred to new bottles right after the hatching (Ashburner, Golic, & Hawley, 2005).

To obtain hybrid males we intercrossed virgin females of one species with virgin males of heterospecific sister species (7 days aged virgin flies as parents) in no choice mating combination, while the combination of both conspecific and heterospecific males with a single female failed to produce any hybrids as the female always chose conspecific males. When progeny emerged, males and females were sorted and kept in separate vials. To study the development of spermatogenesis 50 matured male hybrids (7-10 days old) were randomly selected for observation. The different mating regimes along with its abbreviations is represented in the Suppplementary table S9.

### Mating latency and copulation duration (Courtship behaviour)

Virgins were isolated in the pupal stage from the respective parental and hybrid progeny and were aged for 7 days, with the help of an aspirator a single male and a female were introduced into an empty vial. Sixteen different combinations were used with both males and females of pure species and F1 hybrid males and females. The pair was observed and courtship time and the duration of copulation were recorded (Spieth, 1974; Coyne & Orr, 2004). The procedure was repeated for 30 pairs (N). The time elapsed until mating from the point where courtship behaviour started was taken as mating latency. Duration of copulation was considered between the time of initiation of copulation to termination. The male was taken out from the observation vial after the completion of the copulation.

### Light and Phase contrast microscopy

To investigate the developmental stages of spermatids and assess sperm motility in both parental and hybrid male flies, testes together with seminal vesicles were carefully dissected away from the accessory glands and external genitalia. The tissues were mounted in a drop of phosphate-buffered saline (PBS) and gently squashed under a coverslip, following the protocol of Kemphues et al. (1980). Preparations were examined using a phase-contrast light microscope, and representative images were captured at 40× magnification (Sitaram, Hainline, & Lee, 2014).

### Female sperm transfer

Reproductive tracts of hybrid females were checked for the presence of sperms within one hour of copulation by dissecting the spermatheca, seminal receptacle, and uterus in 1 X PBS using fine insect needles. The cells were stained using DAPI and observed under Olympus CX 21 microscope (20-40X magnification) for the presence of sperms in any of the reproductive tract organs (Heifetz, Lung, Frongillo, & Wolfner, 2000).

### Baseline mating success

Baseline success was quantified under unmanipulated conditions by recording the proportion of successful copulations in both parental crosses (B, M, P, PA) and heterospecific crosses (MB, PB, PAB, BM, PM, PAM, BP, MP, PAP, BPA, MPA, PPA). Each cross was tested with 30 pairs, and success was expressed as mean ± SE across replicates (Refer supplementary table S9, for abbreviations).

### Olfactory manipulation: antennae ablation

To assess the role of olfaction, the antennae were removed following the protocol of Kurtovic, Widmer, & Dickson, 2007). Flies were anesthetized on ice and gently immobilized in truncated pipette tips. Antennae were removed with a sterile razor blade, taking care to remove both funiculus and arista. Control flies were arista-ablated only, as the arista cannot be removed independently from the antenna. Three experimental groups were tested: male-only ablated, female-only ablated, and both sexes ablated, alongside controls. Ablated flies were allowed to recover for 48 h on fresh food before mating assays.

### Gustatory manipulation: foreleg tarsi ablation

To test the role of gustation, foreleg tarsi were removed following Montell (2009). Flies were anesthetized and the five tarsal segments of both forelegs were excised with micro-scissors under a stereomicroscope. Control flies were handled and anesthetized similarly but without tarsal removal. Three groups were tested: male-only ablated, female-only ablated, and both sexes ablated, alongside controls. Ablated flies were allowed to recover for 48 h before mating assays.

### Data analysis

One-way analysis of variance (ANOVA) was performed, keeping the nature of the cross (Conspecific or species hybrids mating) factors. When significant differences were found among the factors, Tukey’s Post hoc multiple comparison tests were performed to test which factor’s averages were significantly different from one another. All statistical tests were conducted in GraphPad Prism 9.5.0. To assess for the overall variation in reproductive traits for both the pure species and hybrids, we performed principal component analysis (PCA) using the prcomp function in R v4.3.2 (R Core Team, 2023). The analysis included three quantitative reproductive measures: copulation duration (minutes), mating latency (minutes), and fertility status (binary coding: 1 = fertile, 0 = sterile). All the variables were centered and scaled prior to the analysis in order to account for the differences in measurement scales, later the first two principal components (PC1 and PC2) were extracted to visualize clustering patterns among species and hybrids. Data visualization was carried out using the ggplot2 package (Wickham, 2016), and 95% confidence ellipses were drawn around groups to highlight separation between parental species and hybrids. Remaining statistical analyses were performed in Python 3.10, with significance set at p < 0.05 (Wickham, 2016; R Core Team, 2023; GraphPad Software, 2022).

## RESULTS

### Mating latency and Copulation duration of pure and hybrid crosses

Heterospecific matings from crosses between *Drosophila bipectinata* sister species complex are fully sterile as the hybrid males fail to produce any progeny either with hybrid females or even when backcrossed with parental females. Nevertheless, it is unrevealed whether sterile males suffer from any kind of reduced pre-copulatory reproductive fitness when compared to fertile parental males. Mating latency and copulation duration were measured for the interspecies hybrid males and also for the parental species. There was a significant difference in mating latency and copulation duration among parental species and hybrid males crossed with hybrid females. One-way ANOVA and Tukey’s multiple comparison test showed significant variation for mean duration of mating latency (F=65.87; df=15, 412; P=<0.0001) in all 16 different pure and hybrid crosses. It is observed that maximum mating latency was observed in hybrid crosses of *D. pseudoananassae* X *D. bipectinata* (F1PAB, n=26), *D. pseudoananassae* X *D. malerkotliana*(F1PAM, n=25), *D. pseudoananassae* X *D. parabipectinata* (F1PAP, n=24), *D. bipectinata* X *D. pseudoananassae*(F1BPA, n=24) and *D. malerkotliana* X *D. pseudoananassae*(F1MPA, n=24) (Fig. 1). Conspecific matings of parental species of *bipectinata* species complex was found to be having lesser duration of mating latency compared to hybrid matings.

**Fig. 1:**
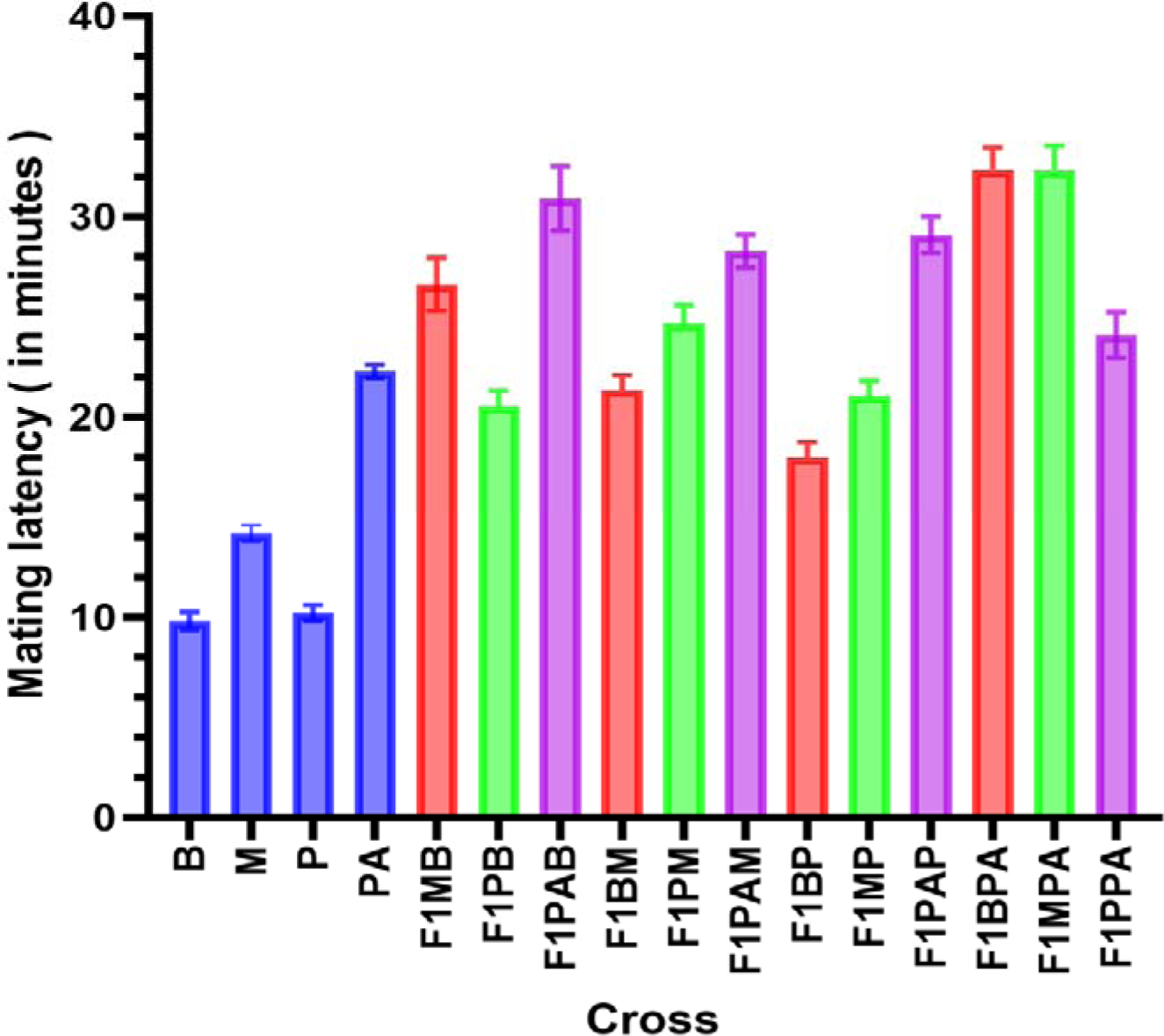
Duration (mins.) of Mating latency in pure and species hybrids of *Drosophila bipectinata* species complex (For labels refer to Supplementary table S9). (F=65.87; df=15, 412; P=<0.0001). Error bars represent +/− 1 standard error.

Tukey’s Post-Hoc test determines that F1 hybrid males (F_1BM,_ n=26; F_1PM,_ n=29 and F_1BP,_ n=28; F_1MP,_ n=26) mated for the same average amount of time as *D. malerkotliana*, n=33 (F=220.9; df=3, 106; P=<0.001) and *D. parabipectinata* parents, n=33 (F=76.27; df=3, 106; P=<0.001) (Fig. 2 and 3), also F_1MB_ (n=26) and F_1PB_ (n=28) hybrid males showed the almost same average amount of copulation duration as *D. bipectinata* parents, n=33 (F=470.2; df=3, 106; P=<0.001). Although we found a non-significant difference among the mating pairs of *D. malerkotliana* vs F_1BM,_ F_1PM_ and *D. parabipectinata* vs F_1BP,_ F_1MP,_ there is a significant difference in the copulation duration of sterile hybrid males based on which of the female parent they have inherited from (Fig. 2,3 and 4).

**Fig. 2:**
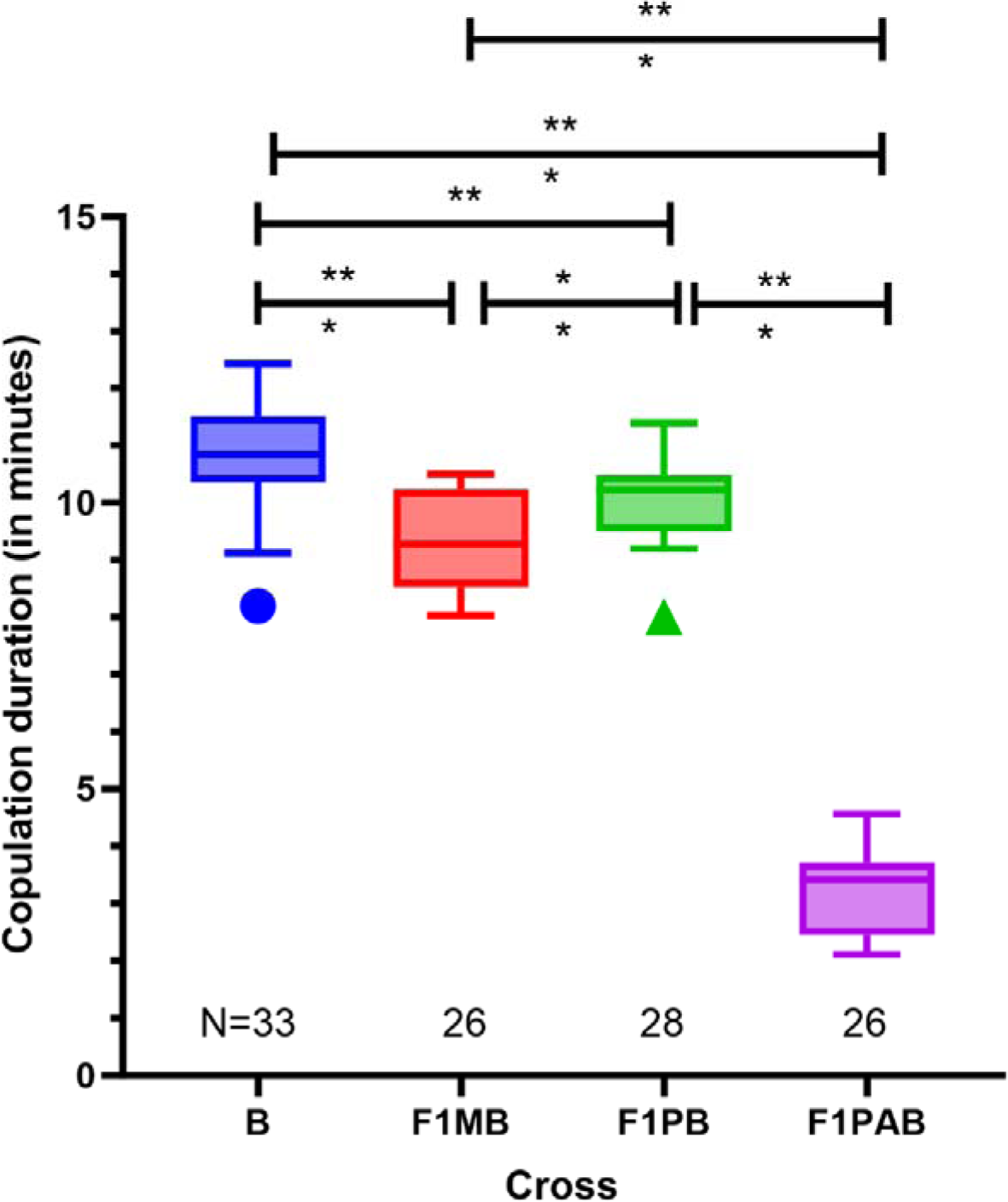
Copulation duration (mins.) assays of *Drosophila bipectinata* pure and species hybrids represented in boxplot (F=470.2; df=3, 106; P=<0.001). Lower boundary of the box indicates 25^th^ percentile, upper boundary of the box indicates 75^th^ percentile of copulation duration in minutes, line inside the box marks the median. Upper and lower whiskers indicate the 10^th^ and 90^th^ percentiles, and icons below the whiskers represent the outliers falling outside the 10^th^ percentile. Bars (Black) above the boxplot represent Tukey’s multiple comparisons between the conspecific and hybrid mating groups and asterisks indicate the significance level (**<0.01, *** <0.001, **** <0.0001). B = *D. bipectinata* females mated to conspecific males (N=33). F1MB(Hybrids from *D. malerkotliana* male and *D. bipectinata* female parents, N=26), F1PB(Hybrids from *D. parabipectinata* male and *D. bipectinata* female parents, N=28), and F1PAB (Hybrids from *D. pseudoananassae* male and *D. bipectinata* female parents, N=26). N = Number of replicates tested.

**Fig. 3:**
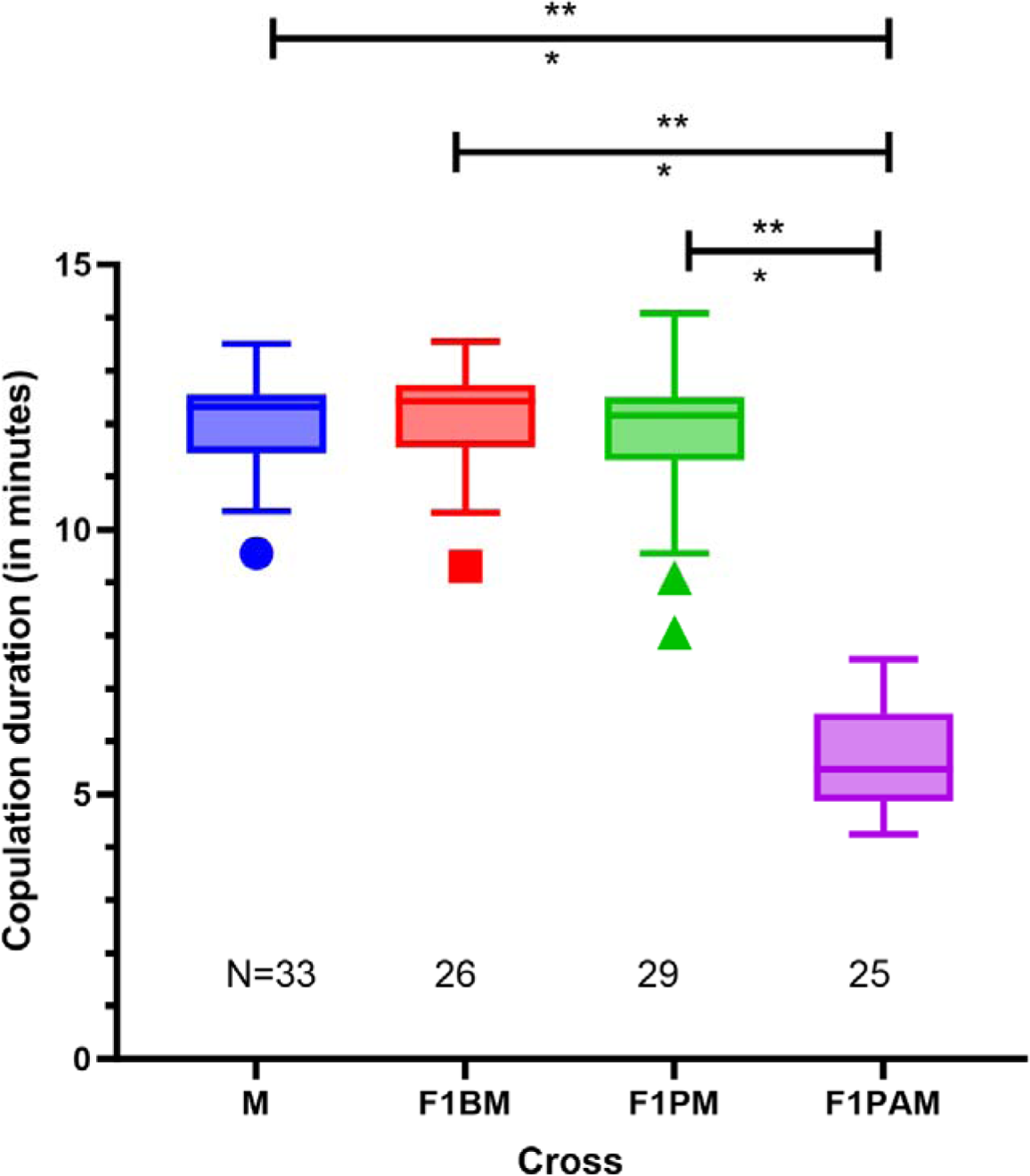
Copulation duration (mins.) assays of *Drosophila malerkotliana* pure and species hybrids represented in boxplot (F=220.9; df=3, 106; P=<0.001). Lower boundary of the box indicates 25^th^ percentile, upper boundary of the box indicates 75^th^ percentile of copulation duration in minutes, line inside the box marks the median. Upper and lower whiskers indicate the 10^th^ and 90^th^ percentiles, and icons below the whiskers represent the outliers falling outside the 10^th^ percentile. Bars (Black) above the boxplot represent Tukey’s multiple comparisons between the conspecific and hybrid mating groups and asterisks indicate the significance level (**<0.01, *** <0.001, **** <0.0001). M = *D. malerkotliana* females mated to conspecific males (N=33). F1BM(Hybrids from *D. bipectinata* male and *D. malerkotliana* female parents, N=26), F1PM(Hybrids from *D. parabipectinata* male and *D. malerkotliana* female parents, N=29), and F1PAM(Hybrids from *D. pseudoananassae* male and *D. malerkotliana* female parents, N=25). N = Number of replicates tested.

The sterile F_1_ hybrid progeny of *D. pseudoannannassae* parents, irrespective of male or female, did not correlate with the copulation duration of female parents (F=364.5; df=3, 96; P=<0.0001) (Fig. 5). The F1 hybrids males (F_1BM_, F_1PM_ and F_1BP,_ F_1MP_) mated to F1 hybrid females grouped with *D. malerkotliana* and *D. parabipectinata* parents, where the copulation duration of these hybrid males is similar to that of maternal parent used to produce particular hybrids, also the F_1MB_ and F_1PB_ hybrid males show the same pattern, almost similar to the *D. bipectinata* copulation duration which implicates that the duration of copulation could be strongly influenced by the X-chromosome of hybrid genome inheritance.

**Fig. 4:**
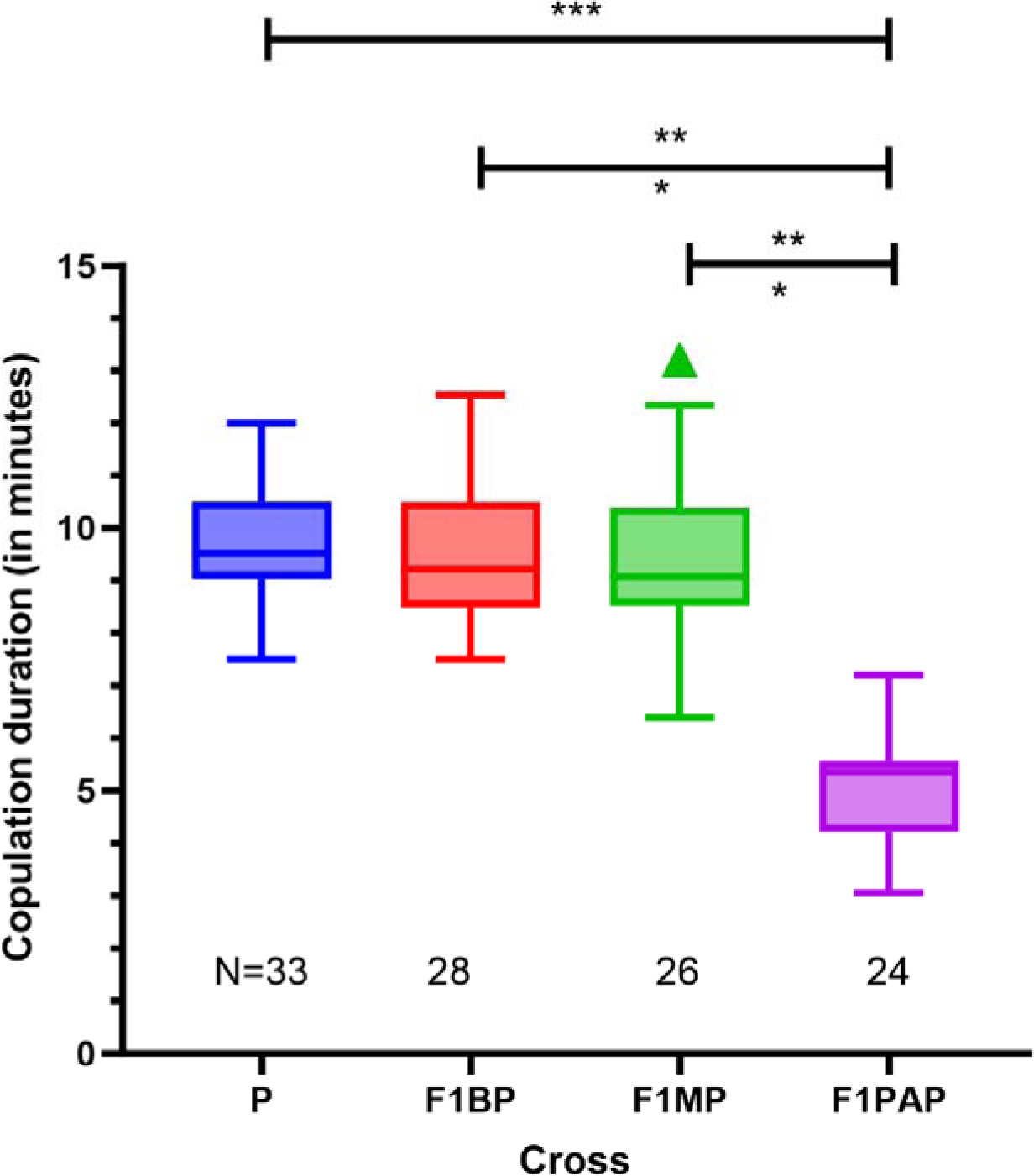
Copulation duration (mins.) assays of *Drosophila parabipectinata* and species hybrids represented in boxplot (F=76.27; df=3, 104; P=<0.001). Lower boundary of the box indicates 25^th^ percentile, upper boundary of the box indicates 75^th^ percentile of copulation duration in minutes, line inside the box marks the median. Upper and lower whiskers indicate the 10^th^ and 90^th^ percentiles, and icons below the whiskers represent the outliers falling outside the 10^th^ percentile. Bars (Black) above the boxplot represent Tukey’s multiple comparisons between the conspecific and hybrid mating groups and asterisks indicate the significance level (**<0.01, *** <0.001, **** <0.0001). P = *D. parabipectinata* females mated to conspecific males (N=33). F1BP(Hybrids from *D. bipectinata* male and *D. parabipectinata* female parents, N=28), F1MP(Hybrids from *D. malerkotliana* male and *D. parabipectinata* female parents, N=26), and F1PAP(Hybrids from *D. pseudoananassae* male and *D. parabipectinata* female parents, N=24). N = Number of replicates tested.

**Fig. 5:**
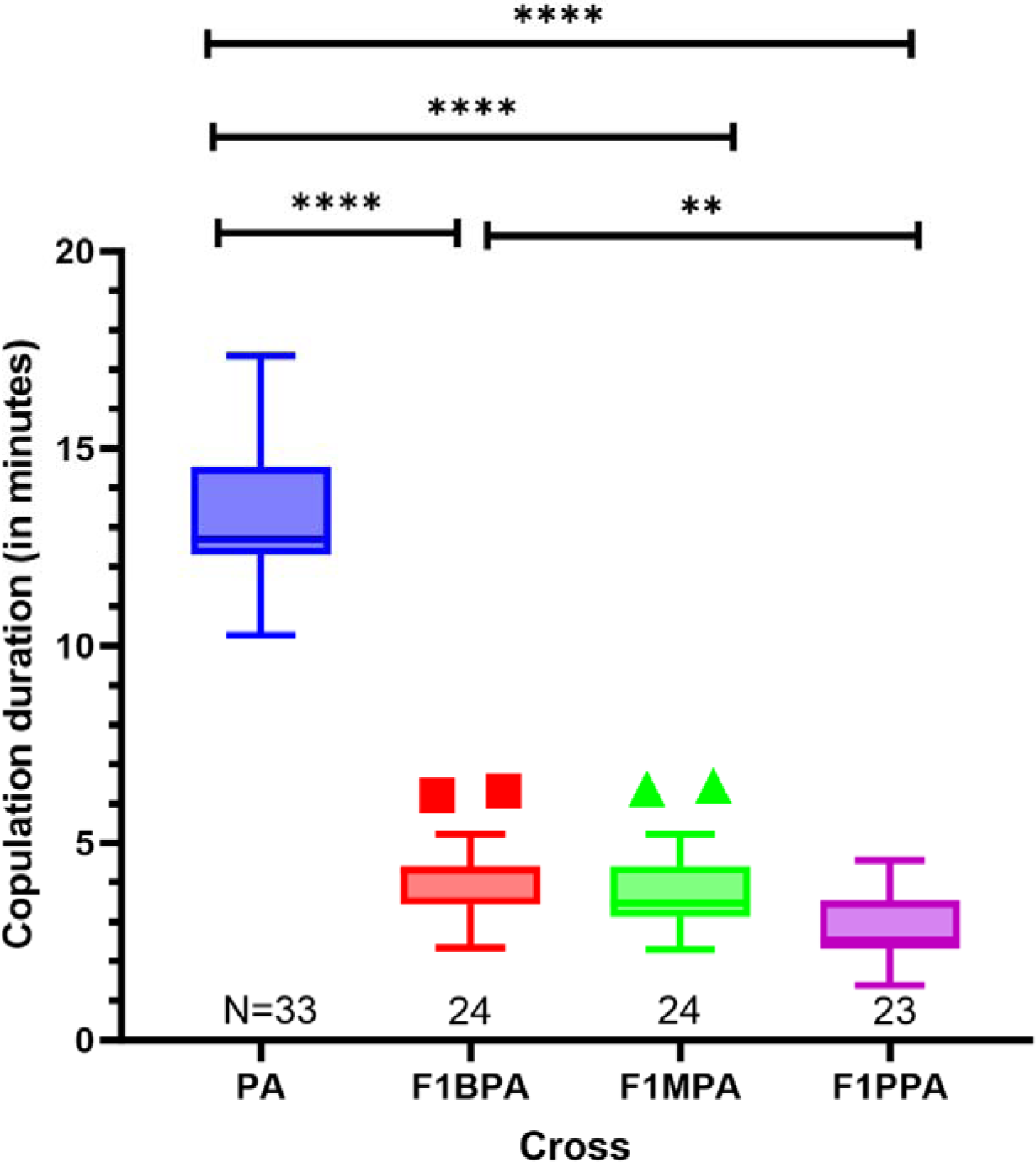
Copulation duration (mins.) assays of *Drosophila pseudoananassae* and species hybrids represented in boxplot(F=364.5; df=3, 96; P=<0.0001). Lower boundary of the box indicates 25^th^ percentile, upper boundary of the box indicates 75^th^ percentile of copulation duration in minutes, line inside the box marks the median. Upper and lower whiskers indicate the 10^th^ and 90^th^ percentiles, and icons below the whiskers represent the outliers falling outside the 10^th^ percentile. Bars (Black) above the boxplot represent Tukey’s multiple comparisons between the conspecific and hybrid mating groups and asterisks indicate the significance level (**<0.01, *** <0.001, **** <0.0001). PA = *D. pseudoananassae* females mated to conspecific males (N=33). F1BPA(Hybrids from *D. bipectinata* male and *D. pseudoananassae* female parents, N=24), F1MPA(Hybrids from *D. malerkotliana* male and *D. pseudoananassae* female parents, N=24), and F1PPA(Hybrids from *D. parabipectinata* male and *D. pseudoananassae* female parents, N=23). N = Number of replicates tested.

### Hybrid males fail to transfer sperm

Even though some of the F1 hybrid males showed a significant reduction in copulation duration compared to their parents, there is a possibility of sperm transfer in that short period of copulation. After the copulation, we separated the female hybrid flies into separate vials and checked for the presence of eggs laid from all the F1 hybrid females for 15 consecutive days, surprisingly we found few number of eggs laid in all the vials, but none of the eggs were fertile and did not show any presence of larvae or pupae, which indirectly concludes that F1 hybrid males either have failed to transfer sperms or there is a possibility of sperm spillage or sperm dumping from the F1 hybrid females. Ovulation in *Drosophila* females is mainly triggered by some of the male factors during and after mating, such as sperm, and mainly by some of the seminal fluid proteins. It is found in *D. melanogaster*, females produces oocytes and store them in an arrested and inactivated state inside the ovary until the copulation happens with a mate, just like many other insects (Wolfner et al. 2003) and egg activation is independent of the sperm entry (Doane 1960, Went and Krause 1974, Macdonald and Struhl 1986, Page & Orr-Weaver 1997). It is also known that one of the seminal fluid protein, Acp26a (ovulin) stimulates ovulation and mediate oocytes release(Heifetz et al. 2000, Heifetz et al. 2001) and there is another example of male ejaculatory duct peptide Dup99B (Fan et al. 2000, Saudan et al. 2002), known to stimulate egg production in unmated flies. So, even if the hybrid males have failed to transfer sperms, the transfer of seminal proteins and ejaculatory duct ovulation-stimulating substances could have triggered the maturation of oocytes even without successful fertilization among the female hybrids.

### Reproductive tracts of hybrid males and females

We conducted the F1 hybrid matings again by producing the hybrids through parental heterospecific matings and assayed the sperm storage organ (spermatheca) of the F1 female hybrids(n=36, 3 hybrid females from each heterospecific matings) for the presence of sperms within one hour after copulation. But none of the female hybrids showed any sperms inside the spermatheca and seminal receptacle, which specifies that hybrid males have failed to transfer sperms or there is a possibility of faster sperm spillage by the hybrid females within 30 minutes after copulation. Hence in order to check the motility of sperms in hybrid males testes along with seminal vesicles were observed through phase contrast microscopy, motile sperms were seen in the 6 days old testes of pure species respectively (Supplementary Figure 1), but hybrid males failed to produce motile sperms. (Supplementary Figure 1-3).

### PCA of reproductive traits

We performed a principal component analysis (PCA) using three reproductive traits: copulation duration, mating latency, and fertility status across parental species and F1 hybrids. The first two principal components explained over 92% of the total variance (PC1 = 76.1%; PC2 = 16.5%). Parental species (*D. bipectinata, D. malerkotliana, D. parabipectinata, D. pseudoananassae*) formed a compact cluster on the right side of the PCA plot, reflecting consistent reproductive performance within pure species. By contrast, hybrids were clearly displaced into separate regions of PCA space. Among hybrids, F1MB and F1PB crosses clustered in the central-upper quadrant, indicating intermediate reproductive values driven by prolonged mating latency. F1PAB and F1PPA crosses, in contrast, were shifted to the lower-left quadrant, reflecting markedly reduced copulation duration combined with extreme sterility (Fig. 6). This separation within hybrids highlights that although sterility is universal, the behavioral reproductive traits (mating latency and copulation duration) vary depending on the parental combination and maternal species.

**Fig. 6.**
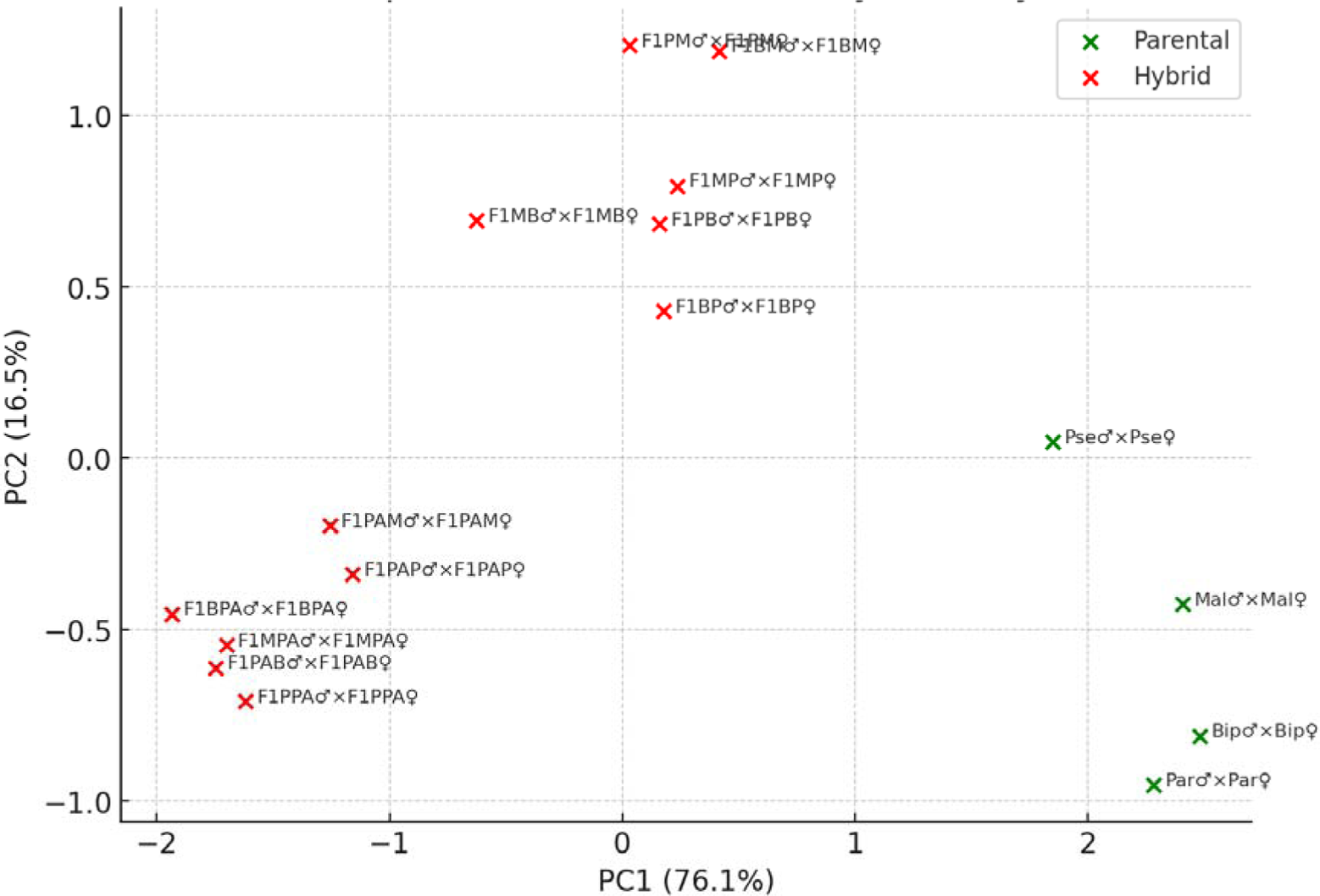
Principal component analysis (PCA) of reproductive traits in parental and hybrid crosses of the *Drosophila bipectinata* complex.

PCA was performed using copulation duration, mating latency, and fertility status across four parental species (*D. bipectinata*, *D. malerkotliana*, *D. parabipectinata*, and *D. pseudoananassae*) and their FLJ hybrids. The first two principal components explain 92.6% of the total variance (PC1 = 76.1%; PC2 = 16.5%). Parental species (green) cluster tightly on the right side of the plot, reflecting uniform reproductive performance, whereas hybrids (red) are displaced into distinct regions of trait space. F1MB and F1PB hybrids group in the central-upper quadrant, showing prolonged mating latency, while F1PAB and F1PPA hybrids shift to the lower-left quadrant, with reduced copulation duration and extreme sterility. These patterns reveal cross-specific combinations of reproductive disruptions underlying hybrid male sterility.

### Baseline heterospecific mating success under control conditions

Under control (unmanipulated) conditions,i.e, without any olfactory or gustatory abberations, heterospecific copulation success rate was consistently low across all the species pairs of *D. bipectinata* complex (Fig. 7). The average proportion of successful copulations ranged from 8–20%, with no single heterospecific mating exceeding more than 20%. This consistent low baseline in the success rate indicates a strong prezygotic isolatory mechanisms reinforcing the barrier between the species, even in the absence of experimental manipulations. Naturally intraspecific parental crosses showed significantly higher success rates (≥75%), forming the reference point against the sensory manipulations evaluation.

**Fig. 7:**
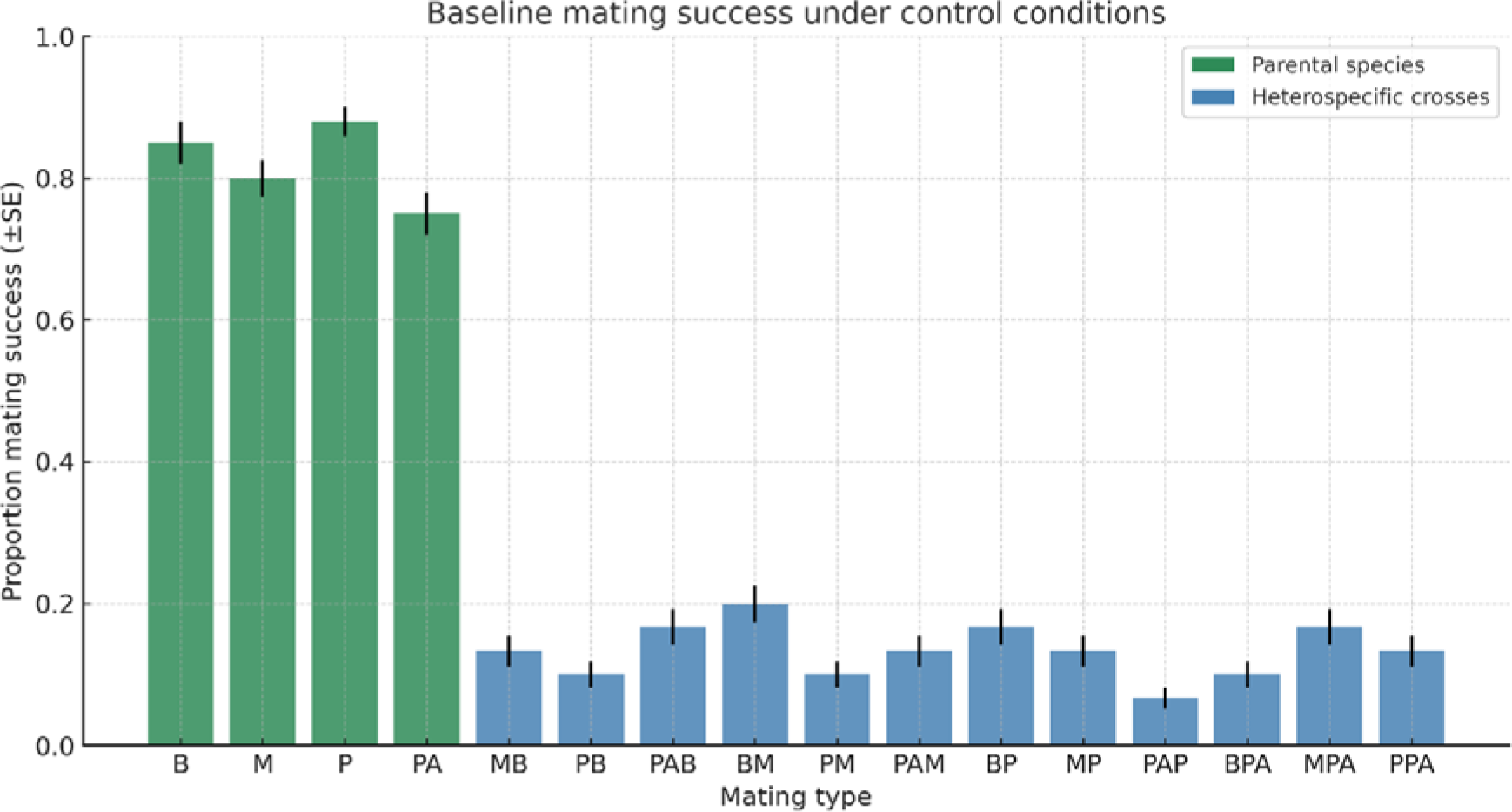
Baseline heterospecific mating success under control conditions. Bar plot showing the proportion of successful copulations (±SE) for all heterospecific crosses in the *D. bipectinata* species complex. Success rates were generally low (≤20%), confirming strong prezygotic isolation under unmanipulated conditions.

### Effect of olfactory manipulations (antennae ablation)

To assess the role of olfactory input in mating success, antennae were ablated in either males, females, or both sexes, with aristae-removed flies serving as controls. Antennal ablation in males caused a significant reduction in copulation success across all heterospecific crosses (average 4–7% success vs. 12–18% in controls, p < 0.001; Fig. 8A,B). Female ablation produced milder effects (average 8–12%), while ablation in both sexes almost completely abolished mating (1–3%). These results highlight that male olfactory input is indispensable for successful heterospecific mating (Fig. 9).

**Fig. 8:**
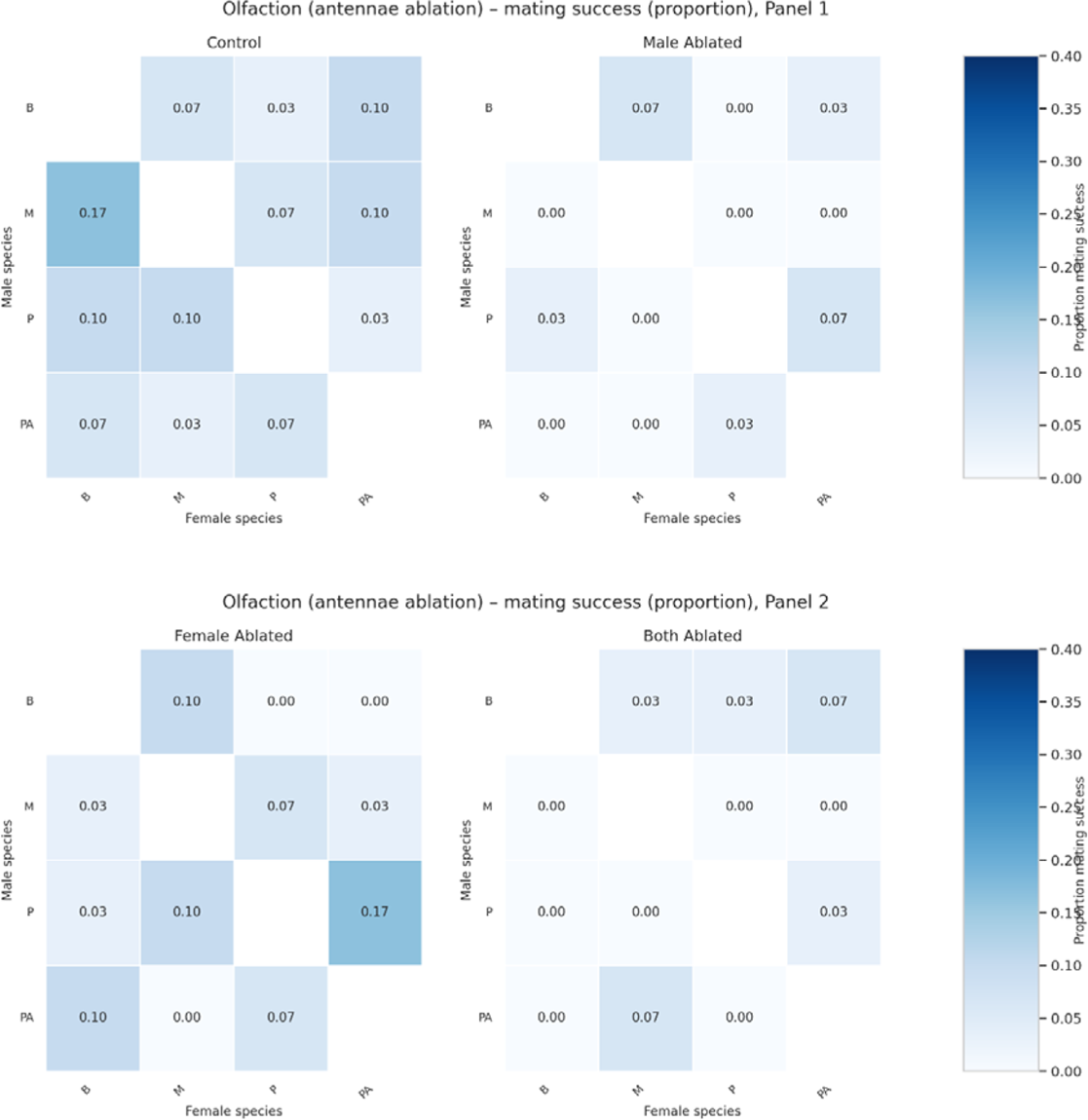
Effect of antennal ablation (olfaction) on heterospecific mating.

**Fig. 9:**
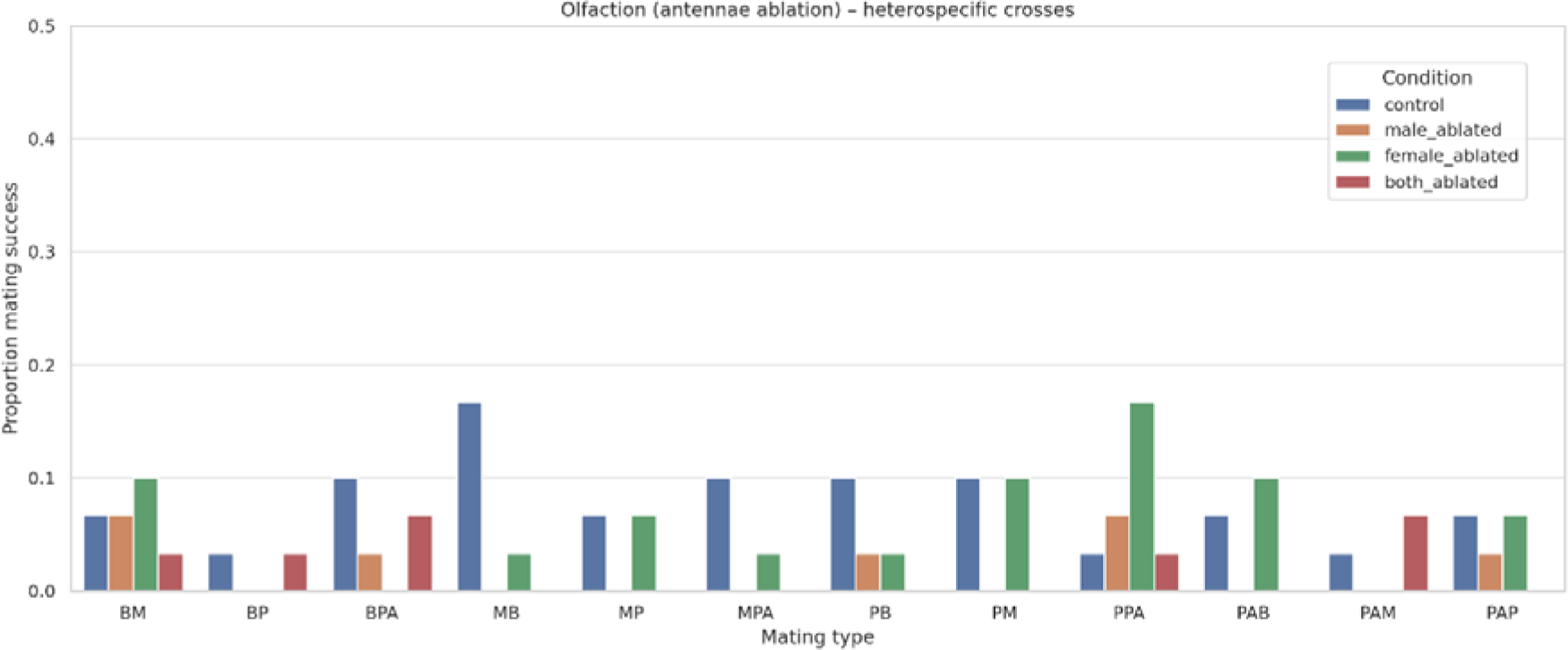
Bar plots (±SE) of heterospecific mating success for each cross (abbreviated). Internal legend indicates experimental conditions. Male antennal removal strongly reduced heterospecific mating; female ablation had milder effects; ablation of both sexes nearly abolished copulation.

Heatmaps (Panel 1: Control, Male antennal ablated; Panel 2: Female antennal ablated, Both ablated) showing the proportion of successful copulations for each cross. Axes are labeled with species abbreviations: B = *D. bipectinata*, M = *D. malerkotliana*, P = *D. parabipectinata*, PA = *D. pseudoananassae*. Color scale (right) represents the proportion of successful matings. Male antennal removal drastically reduced heterospecific mating; female ablation had milder effects; ablation of both sexes nearly abolished copulation.

### Effect of gustatory manipulations (foreleg tarsi ablation)

Foreleg tarsi removal produced striking sex-specific outcomes. Male tarsi ablation significantly increased heterospecific mating success (average 22–28% vs. 12–18% in controls, p < 0.01), suggesting that male gustatory receptors normally act as inhibitory filters that restrict heterospecific courtship. In contrast, female tarsi ablation completely abolished mating success across all crosses, underscoring the requirement of female gustatory inputs for acceptance of copulation (Fig. 10A,B). Ablation of both sexes resulted in near-complete loss of mating (1–4%) (Fig. 11). These results establish gustation as a primary determinant of female mate choice and a secondary filter in males.

**Fig. 10:**
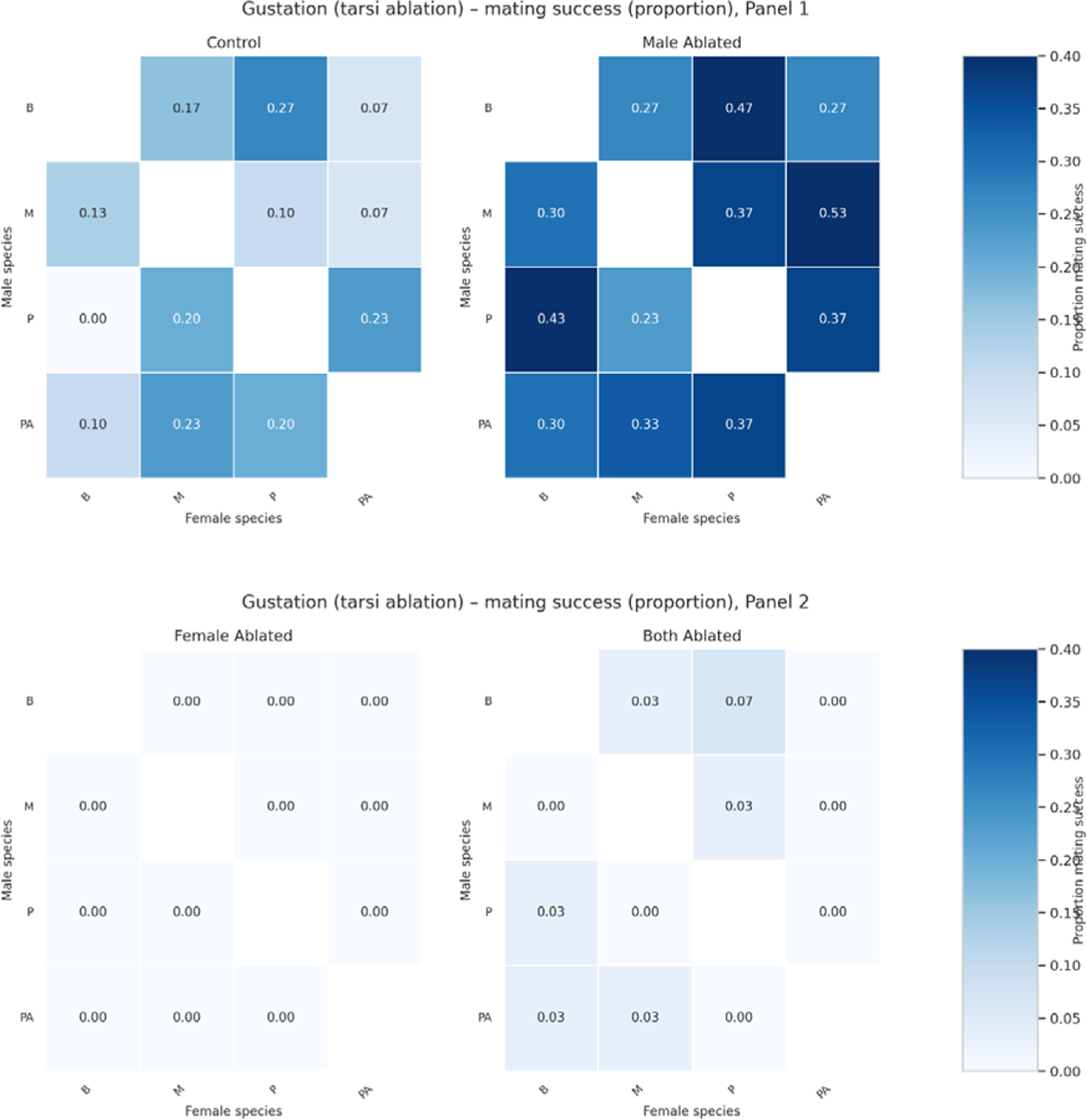
Effect of foreleg tarsi ablation (gustation) on heterospecific mating.

**Fig. 11:**
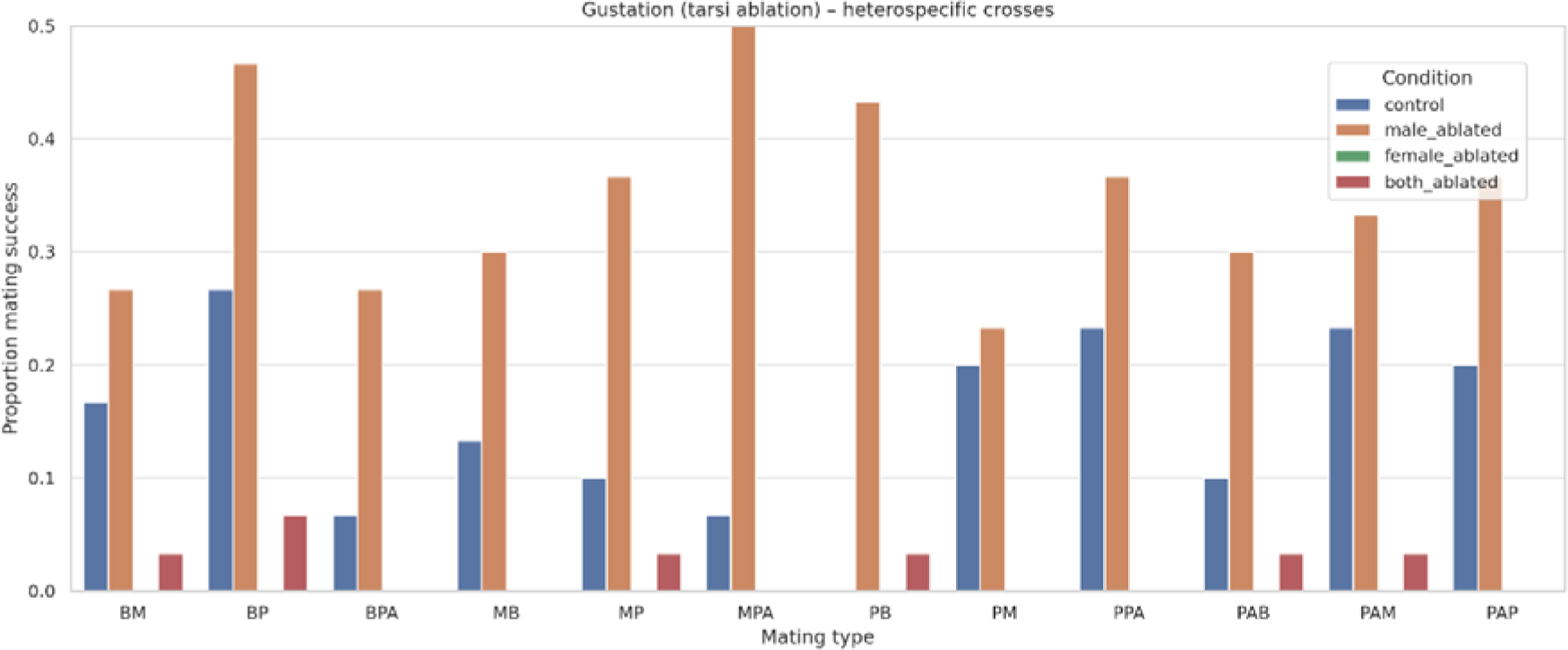
Bar plots (±SE) of heterospecific mating success for each cross (abbreviated). Internal legend indicates experimental conditions. Male tarsi removal markedly increased heterospecific mating, while female tarsi removal abolished it; ablation of both sexes reduced success to near zero.

Heatmaps (Panel 1: Control, Male tarsi ablated; Panel 2: Female tarsi ablated, Both ablated) showing the proportion of successful copulations for each cross. Axes are labeled with species abbreviations: B = *D. bipectinata*, M = *D. malerkotliana*, P = *D. parabipectinata*, PA = *D. pseudoananassae*. Color scale (right) represents the proportion of successful matings. Male tarsi removal increased heterospecific copulations, whereas female tarsi removal abolished them; both ablated reduced success to near zero.

### Comparative effects of olfactory and gustatory manipulations

Comparative analyses showes that antennal ablation consistently reduced heterospecific mating success, but ablation of tarsi, produced sex-specific effects (Fig. 12), which demonstrates that male olfactory inputs were critical in detecting pheromone blends, while female gustatory input were essential in confirming mate identity. Together, these manipulations reveal a dual-sensory system that integrates both olfactory and gustatory modalities to regulate reproductive isolation.

**Fig. 12:**
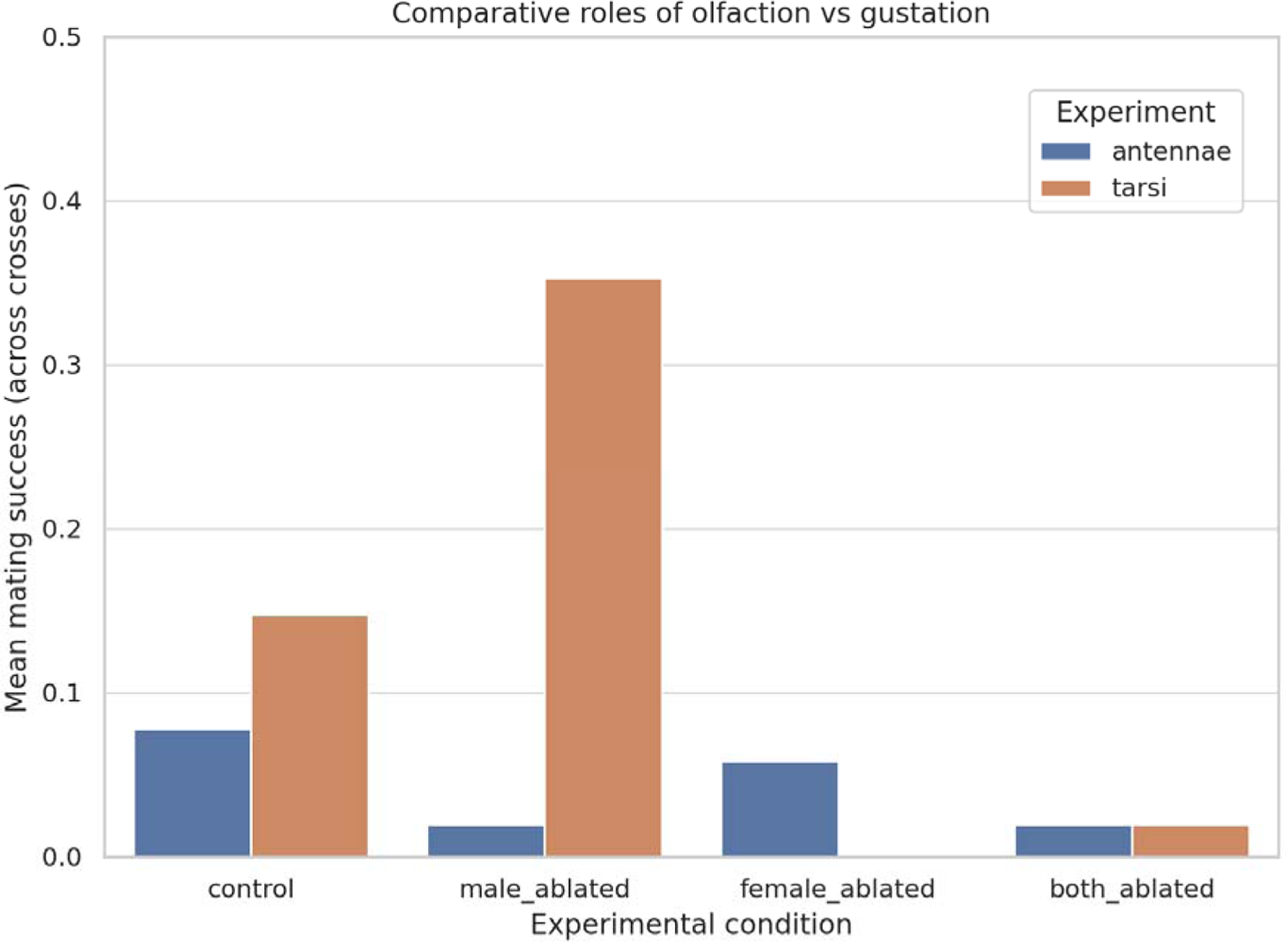
Comparative roles of olfactory and gustatory cues in heterospecific mating.

Bar plot showing mean copulation success (±SE) across all crosses for antennal (olfaction) and tarsi (gustation) manipulations. X-axis indicates experimental conditions (control, male ablated, female ablated, both ablated). Internal legend distinguishes experiments. Olfactory manipulations consistently suppressed heterospecific mating, whereas gustatory manipulations produced sex-specific effects: male ablation increased copulation success, while female ablation abolished it.

## DISCUSSION

The advent of hybrid male sterility in the evolution of genetically diverging species was evident enough for researchers, to state that sterility of heterogametic male hybrids is the first milestone in the process of speciation (Coyne and Orr 1989). Primarily, we observed the latency duration among the parental and hybrid crosses (Fig. 1) through no-choice mating experiments, there were no significant differences among the pure species except for the *D. pseudoananassae* parental cross which took a comparatively longer duration of latency period. All the F1 hybrid crosses with their respective hybrid mates showed significantly longer mating latency which specifies the attractiveness of the males to their females (more the attraction, shorter the latency duration). The longer duration of latency period of the hybrid flies compared to pure species can be attributed to changes in the profile of pheromones in which the male and female specific cuticular hydrocarbons differences among the hybrids could be leading into the delayed female response to the male courtship. Courtship is also one of the innate behaviour consisting of multiple sensory modalities under the strong influence of somatic sex determination hierarchy (Ellis and Carney 2010). Since these hybrids carry genomic backgrounds from considerably divergent sister species, the neural circuitries responsible for courtship behaviour and the genetic makeup regulating the sensory cues and cuticular hydrocarbons could be the reason for the delayed latency period among the hybrids.

Patterns of mating latency and copulation duration themselves provide strong behavioural evidence for reinforcement of reproductive isolation. In our study, the four parental species of the bipectinata complex showed relatively short and consistent mating latencies, with the exception of *D. pseudoananassae*, while all hybrid crosses exhibited significantly prolonged latencies. These delays in the initiation of copulation and reduced copulatory duration are due to divergence in mate recognition systems, such as olfactory or gustatory cues (Coyne et al., 1994; Yukilevich, 2012), since copulation duration was stable between the parental pure species but highly variable and shortened among the hybrids, a pattern which could be associated with the incomplete transfer of seminal proteins leading to reduced reproductive success (Price et al., 2001; Manier et al., 2013). These kinds of disruptions in fundamental behavioural traits suggests that hybrids are doubly disadvantageous: primarily by lower attractiveness and delayed mating, and further by reduced efficiency of copulation once it occurs. Comparable results have been reported in the *D. simulans* clade, where hybrids show longer latency and aberrant copulation behaviour that contribute directly to postzygotic sterility (Wu et al., 1992; Moehring et al., 2007). Thus, even without invoking sensory manipulations, divergence in mating latency and copulation duration between parental species and their hybrids provides clear evidence that behavioural isolation and hybrid dysfunction reinforce one another, strengthening reproductive barriers and promoting speciation

Our experiments demonstrate that both olfactory and gustatory cues are central to the maintenance of reproductive isolation in the *Drosophila* bipectinata species complex, but with distinct sex-specific functions. Olfactory input, particularly from the male antennae, was indispensable for initiating courtship and successful heterospecific copulation. This aligns with earlier studies showing that olfactory cues, mediated via antennal sensilla, guide male orientation and courtship toward conspecific females (Coyne et al., 1994; Kurtovic et al., 2007; Lebreton et al., 2017).

The importance of olfactory cues in the mating discrimination has been closely linked to pheromone detection. In *Drosophila*, the one of the volatile pheromone cis-vaccenyl acetate (cVA) has been detected by the receptor Or67d which is known to regulate mate recognition and aggregation (Kurtovic et al., 2007). One of the recent work has shown that Or67d in *D. bipectinata* species complex exhibits species-specific sensitivity to cVA, which suggests divergence in pheromone reception among the members of this sub-complex (Sharma et al., 2024), additionally other receptors, such as Or69a, also known to integrate responses for both food odors and sex pheromones, which links ecological niche specialization with mate choice (Lebreton et al., 2017). Also, Or47b olfactory neurons adapted to male cuticular pheromones in order to modulate the female receptivity (Yun et al., 2024). Together, these findings provide a mechanistic explanation for our experimental results, in which male antennal input appears to be indispensable in detecting species-specific pheromone blends, while female antennal input, though less critical, likely contributes to acceptance decisions. Divergence in pheromone production and receptor tuning thus offers a plausible basis for the strong reduction in heterospecific success observed after antennal ablation in the bipectinata complex.

Gustatory manipulations produced strikingly asymmetric outcomes. Male tarsi ablation increased heterospecific copulation success, whereas female tarsi ablation abolished it entirely, with both-sex ablation reducing success to near zero, which is consistent with the role of tarsal gustatory bristles and receptors present in tarsi, for detecting cuticular hydrocarbons (CHCs), which acts as a key regulatory contact pheromones in *Drosophila* (Inoshita et al., 2011; Barmina et al., 2022). Male gustatory receptors, such as Gr33a, usually acts as an inhibitory sensors which prevents escalation of courtship behavior towards heterospecific females through the recognition of species-specific CHC blends (Moon et al., 2009; Inoshita et al., 2011). Ablation of male tarsi could be removing this inhibitory checkpoint, thereby increasing heterospecific mating. In contrast, female tarsi appear to provide a permissive cue, confirming the male identity and enabling the copulation paradigm. Removing these critical female tarsi from the individuals likely prevents recognition of male CHCs, ending up as a complete breakdown of mating success, these findings support the idea that gustatory cues especially in females, are not secondary behavior to olfaction but play as a primary role in mate acceptance, acting as a significant barrier in female choice while restricting heterospecific mating via male contact chemoreception.

The PCA analysis of reproductive traits further accentuate the multifactorial nature of reproductive isolation mechanisms among the species complex. Pure species groups clustered tightly, reflecting stable reproductive traits such as latency, copulation duration, and fertility, consistent with successful reproduction. Hybrids on the other side showed wide-displacement across the PCA space, reflecting multilayered reproductive dysfunction, including prolonged latency, altered copulation duration, and near-complete sterility. These findings emphasizes earlier work among the *D. simulans* clade, where hybrid sterility arose from the disruption of multiple reproductive traits rather than a single factor (Wu et al., 1992; Moehring et al., 2007; Sundararajan & Civetta, 2011). The dispersion among hybrid groups in PCA space suggests that, while sterility is a shared outcome, the underlying behavioral disruptions vary depending on cross direction and parental combination, consistent with maternal effects and cytoplasmic–nuclear incompatibilities (Sawamura, 2000; Presgraves, 2010).

Even though many efforts have gone in deciphering the genetic basis of hybrid male sterility, the clearcut ‘phenotypic’ abnormalities of the hybrid sterility are generally being disregarded. Studies at the molecular level of the spermatogenesis in some of the sterile hybrids have shown misregulation of genes at the later stages of sperm development (Michalak and Noor 2003). There are certain hypotheses explaining the genetic basis of hybrid male sterility such as fast-male and faster X evolution which speculates that the genes involved in gametogenesis evolve rapidly at a higher rate compared to female-biased genes (Haertly et al. 2007), in fast X evolutionary hypothesis hybrid male sterility genes are supposedly augmented in X chromosomes than on autosomes (due to more competent fixation of mutations in the X chromosome) (Charlesworth et al. 1987). Even though these detailed studies on the molecular basis of hybrid sterility are helpful, a direct linkage between sterility phenotypes and undergoing spermatogenesis or fertilization problems lays a precise foundation for a better understanding of the speciation progress. In the present study of *D. bipectinata* hybrids, despite being able to engage in copulation for a considerable duration of time and even considering the eggs laid by hybrid females after copulation, at first glance it appears to be hybrid breakdown but observing the morphology of hybrid testes and its undifferentiated sperms it is revealed that hybrid sterility is due to spermatogenic defects and the isolation is happening at the postmating prezygotic stage itself. The ultrastructural and functional process of spermatogenesis in *D. melanogaster* is well-studied and has been classified into four major stages (germline proliferation, mitosis into meiosis transition, sperm maturation, and individualization). In the present study, most of the testes’ morphological atrophies are relevant to later stages of spermiogenesis i.e, elongation and spermatid individualization. Nine out of twelve F1 hybrid males showed defects that were postmeiotic in nature. Our investigation of sterility referring to premeiotic or postmeiotic is differentiated based on the presence of elongated spermatids at the mid-testes. Even though these hybrids reached the spermatid elongation level there is a possibility of defects occurring at the premeiotic level itself. We can observe a wide scale of atrophies, from primary spermatocyte bundles to aspermic testes. For instance, a substantial reduction in the number of sperm bundles which has failed to individualize, and an increase in sperm bundle size is seen in all the hybrid males (Supplementary Figures 1-3).

Another viewpoint on the copulation duration is, if we take into account the larger effect of the X chromosome on the genotype of sterile F1 hybrid males, we can observe that few hybrids which have inherited their X chromosome from the maternal side show the same value of copulation as their parent. In the preliminary studies conducted by Mishra and Singh(2005) to identify the underlying roles of hybrid male sterility in the bipectinata species complex, they hypothesized that hybrid males of *D. bipectinata* and *D. parabipectinata* are sterile as a consequence of incompatible interactions between X and Y or X and autosomes or due to polygenic interactions, in another study made by Kulathinal and Singh 1998, on *D. mauritiana*, *D. sechelia* and *D. simulans* shows that the X chromosome inherited from *D. mauritiana* withstands the largest effect on hybrid male sterility when hybridized with other species of the clade. Taken together, the integration of sensory manipulations and trait-based PCA demonstrates that reproductive isolation in the *D*. *bipectinata* complex arises from a synergistic interaction of multiple barriers. Prezygotic barriers arbitrated by olfactory (pheromone detection) and gustatory (CHC discrimination) cues could be acting along with postzygotic sterility to prevent gene flow between sister species. The sex-specific asymmetry we observed, in which male and female gustatory inputs having opposite roles, underscores the importance of sexual conflict and female choice in shaping the evolution of isolation mechanisms. Our findings reinforce the view that speciation in *Drosophila* is rarely the product of a single isolating mechanism but emerges through the integration of ecological, behavioral, and genetic divergence evolved into a multilayered isolation framework (Coyne & Orr, 2004; Yukilevich, 2012).

## CONSCLUSION

The current study reveals that reproductive isolation in the *Drosophila bipectinata* species complex is a multifaceted process shaped up by both the behavioral and genetic mechanisms. Divergence in latency and copulation duration between parental species and among the hybrids, coupled with disruptions in pheromone detection and cuticular hydrocarbon recognition, emphasize the importance of prezygotic isolation among the F1 hybrids. On the other hand, postzygotic barriers such as spermatogenic defects and hybrid sterility reinforces the speciation boundaries, corroborating the fact that even when heterospecific matings does occur, they rarely result in viable gene flow. The sex-specific roles of sensory modalities such as olfactory and gustatory cues, together with the disproportionate influence of the X chromosome on hybrid dysfunction, underscores how behavioral, ecological, and molecular factors interact accordingly in reinforcing the species boundaries. Thus, speciation in *D. bipectinata* species complex is best understood as the outcome of a synergistic interplay of multiple reproductive isolating mechanisms acting at different stages of reproduction, giving a comprehensive view of how barriers accumulate and reinforce one another during evolutionary divergence.

## ACKNOWLEDGMENTS

We thank the Department of Studies in Zoology, University of Mysore for providing the infrastructure and We acknowledge the Council of Scientific and Industrial Research (CSIR), Govt. of India for providing NET fellowship.

## AUTHOR CONTRIBUTIONS

Manjunath M: Conceptualization, Formal analysis, Methodology, Investigation, Data curation, Writing—original draft, Software, Visualization.

Damini C S: Writing—review & editing

Dr. Shakunthala V: Project administration, Validation, Resources, Writing—review & editing.

