## supplementary table s9 and figures for "Reinforcement of speciation by behavioural and reproductive barriers in the *Drosophila bipectinata* species complex"

**Supplementary table S9: Crosses involved in the present study-**Abbreviations representing their respective mating types and the hybrid regimes involved in the current study

| **Si. no** | **Abbreviation** | **Mating type** | **Hybrid regime** |
| --- | --- | --- | --- |
| 1 | B | *Drosophila bipectinata*♂ X *Drosophila bipectinata*♀ | - |
| 2 | M | *Drosophila malerkotliana*♂ X *Drosophila malerkotliana*♀ | - |
| 3 | P | *Drosophila parabipectinata*♂ X *Drosophila parabipectinata*♀ | - |
| 4 | PA | *Drosophila pseudoananassae*♂ X *Drosophila pseudoananassae* ♀ | - |
| 5 | F1MB | F1 Hybrid ♂ X F1 Hybrid ♀ | *D. malerkotliana*♂ X *D. bipectinata*♀ |
| 6 | F1PB | F1 Hybrid ♂ X F1 Hybrid ♀ | *D. parabipectinata*♂ X *D. bipectinata*♀ |
| 7 | F1PAB | F1 Hybrid ♂ X F1 Hybrid ♀ | *D. pseudoananassae*♂ X *D. bipectinata*♀ |
| 8 | F1BM | F1 Hybrid ♂ X F1 Hybrid ♀ | *D. bipectinata*♂ X *D. malerkotliana*♀ |
| 9 | F1PM | F1 Hybrid ♂ X F1 Hybrid ♀ | *D. parabipectinata*♂X *D. malerkotliana*♀ |
| 10 | F1PAM | F1 Hybrid ♂ X F1 Hybrid ♀ | *D. pseudoananassae*♂ X *D. malerkotliana*♀ |
| 11 | F1BP | F1 Hybrid ♂ X F1 Hybrid ♀ | *D. bipectinata*♂ X *D.* *parabipectinat*a♀ |
| 12 | F1MP | F1 Hybrid ♂ X F1 Hybrid ♀ | *D. malerkotliana*♂X *D. parabipectinata*♀ |
| 13 | F1PAP | F1 Hybrid ♂ X F1 Hybrid ♀ | *D. pseudoananassae*♂ X *D. parabipectinata*♀ |
| 14 | F1BPA | F1 Hybrid ♂ X F1 Hybrid ♀ | *D. bipectinata*♂ X *D. pseudoananassae* ♀ |
| 15 | F1MPA | F1 Hybrid ♂ X F1 Hybrid ♀ | *D. malerkotliana*♂ X *D. pseudoananassae* ♀ |
| 16 | F1PPA | F1 Hybrid ♂ X F1 Hybrid ♀ | *D. parabipectinata*♂ X *D. pseudoananassae* ♀ |

**Supplementary Figures**

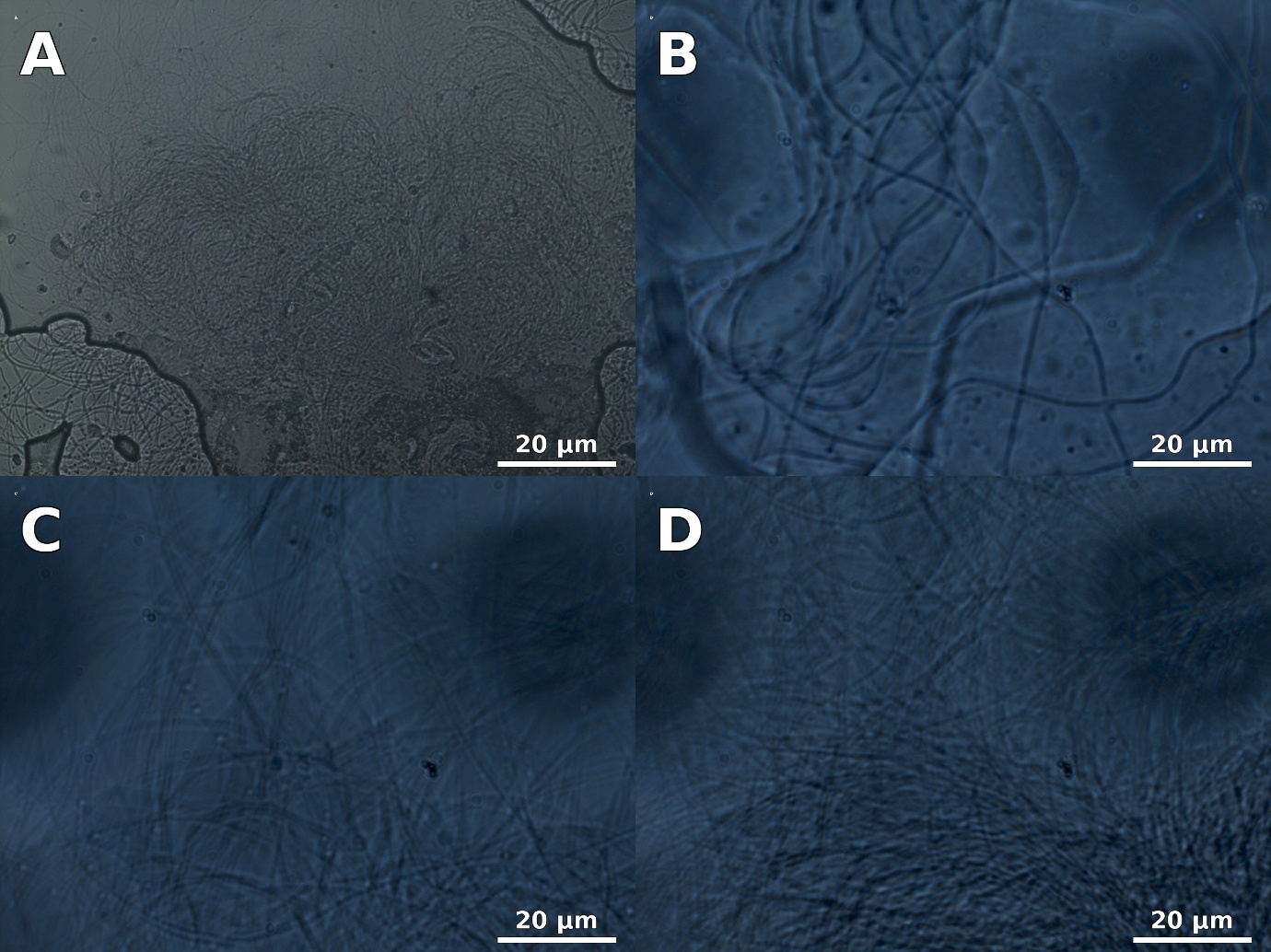

**Supplementary Figure** 1: Phase contrast micrographs of testes squash preparations from 4-day-old parental males of the Drosophila bipectinata species complex.

Panels (A) *D. bipectinata*, (B) *D. malerkotliana*, (C) *D. parabipectinata*, and (D) *D. pseudoananassae* show fully differentiated testes with abundant elongated spermatids and clearly visible individualized sperm bundles (arrows). The presence of individualized sperm indicates normal spermiogenesis and complete differentiation from spermatocytes to motile spermatozoa, reflecting the fertility of pure species. These images provide a baseline against which hybrid defects were compared. (Magnification: 40–100×; Scale bar: 10–20 μm).

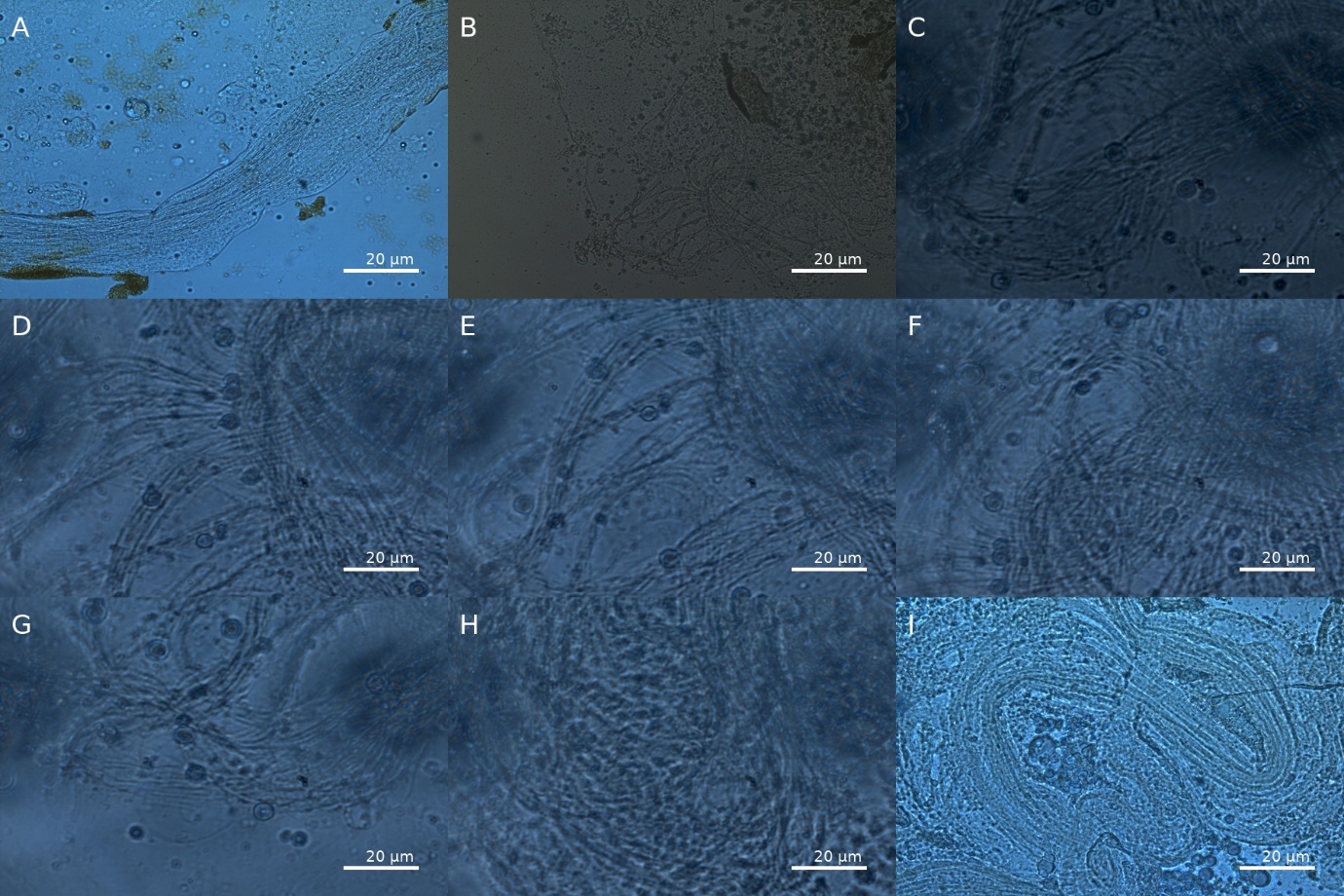

**Supplementary Figure** 2: Phase contrast micrographs of mid-testes squashes from post-meiotically defective hybrid males of the *Drosophila bipectinata* complex.

Hybrid testes exhibit severe defects during the spermiogenesis stage, showing clusters of unindividualized spermatids rather than mature sperm bundles. Panels depict representative crosses: (A) *D. bipectinata*♂ × *D. malerkotliana*♀, (B) *D. bipectinata*♂ × *D. parabipectinata*♀, (C) *D. malerkotliana*♂ × *D. bipectinata*♀, (D) *D. malerkotliana*♂ × *D. parabipectinata*♀, (E) *D. pseudoananassae*♂ × *D. bipectinata*♀, (F) *D. pseudoananassae*♂ × *D. malerkotliana*♀, (G) *D. parabipectinata*♂ × *D. bipectinata*♀, (H) *D. parabipectinata*♂ × *D. malerkotliana*♀, and (I) *D. pseudoananassae*♂ × *D. malerkotliana*♀. In all cases, spermatids fail to undergo proper elongation and individualization, resulting in abnormal sperm bundles and aggregated spermatids. These phenotypes indicate post-meiotic disruption as the underlying mechanism of sterility. (Magnification: 40×; Scale bar: 20 μm).

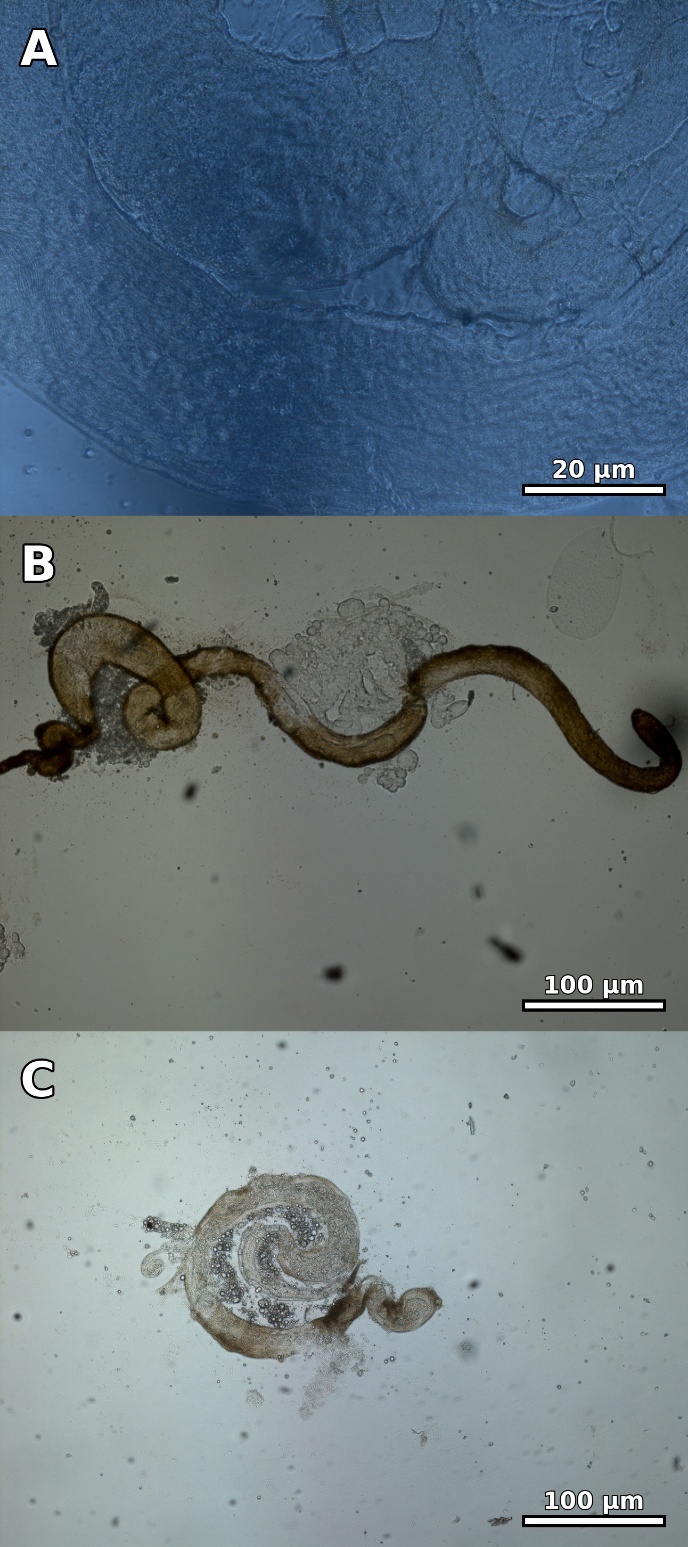

**Supplementary Figure** 3: Fig. 3. Aspermic testes arrested at pre-meiotic stages in sterile hybrids of the *Drosophila bipectinata* complex.

Testes from hybrids (A) *D. bipectinata*♂ × *D. pseudoananassae*♀, (B) *D. malerkotliana*♂ × *D. pseudoananassae*♀, and (C) *D. parabipectinata*♂ × *D. pseudoananassae*♀ reveal severe developmental arrest prior to spermatocyte differentiation. In these crosses, germ cells fail to progress into primary spermatocytes, resulting in aspermic testes with a complete absence of elongating or individualized spermatids. This pattern demonstrates pre-meiotic failure, contrasting with the post-meiotic defects observed in other hybrid combinations (Fig. 2), and highlights the diverse pathways through which hybrid sterility is manifested. (Magnification: 40×; Scale bar: 20-100 μm).
